# Threshold regulation and stochasticity from the MecA/ClpCP proteolytic system in *Streptococcus mutans* competence

**DOI:** 10.1101/263210

**Authors:** M. Son, J. Kaspar, S.J. Ahn, R.A. Burne, S.J. Hagen

## Abstract

Many bacterial species use the MecA/ClpCP proteolytic system to block entry into genetic competence. In *Streptococcus mutans*, MecA/ClpCP degrades ComX (also called SigX), an alternative sigma factor for the *comY* operon and other late competence genes. Although the mechanism of MecA/ClpCP has been studied in multiple *Streptococcus* species, its role within noisy competence pathways is poorly understood. *S. mutans* competence can be triggered by two different peptides, CSP and XIP, but it is not known whether MecA/ClpCP acts similarly for both stimuli, how it affects competence heterogeneity, and how its regulation is overcome. We have studied the effect of MecA/ClpCP on the activation of *comY* in individual *S. mutans* cells. Our data show that MecA/ClpCP is active under both XIP and CSP stimulation, that it provides threshold control of *comY*, and that it adds noise in *comY* expression. Our data agree quantitatively with a model in which MecA/ClpCP prevents adventitious entry into competence by sequestering or intercepting low levels of ComX. Competence is permitted when ComX levels exceed a threshold, but cell-to-cell heterogeneity in MecA levels creates variability in that threshold. Therefore MecA/ClpCP provides a stochastic switch, located downstream of the already noisy *comX*, that enhances phenotypic diversity.

## Introduction

Many species of streptococci can become naturally transformable by entering the transient physiological state known as genetic competence (Fontaine *et al.*, 2014; Johnston *et al.*, 2014). Competence plays a particularly important role for the oral pathogen *Streptococcus mutans*, influencing cell growth, death, interactions with other members of the oral flora and expression of known virulence traits. Bacteriocin production, biofilm formation, acid production and tolerance of acid and oxidative stresses by *S. mutans* all facilitate the competition, persistence and virulence of this organism in the human oral biofilm environment (J. A. Lemos; Burne, 2008). All of these traits are linked to the expression of ComX (also called SigX), an alternative sigma factor that activates competence genes required for DNA uptake and processing. ComX production is controlled by a pathway that integrates signals received from two quorum sensing peptides (Shanker; Federle, 2016) with environmental cues such as pH (Guo *et al.*, 2014; Son *et al.*, 2015b) and oxygen and reactive oxygen species (De Furio *et al.*, 2017), intracellular noise (stochasticity) and positive and negative feedback (Smith; Spatafora, 2012; LeungDufour *et al.*, 2015; Reck *et al.*, 2015; Son *et al.*, 2015a; Hagen; Son, 2017). As a result, *S. mutans* competence is a complex and heterogeneous behavior that can be exquisitely sensitive to the extracellular environment and that remains incompletely understood.

Population heterogeneity in *S. mutans* competence is evident from the low efficiency of natural genetic transformation (Y. Li *et al.*, 2001), as well as from observations of cell-to-cell variability in *comX* gene expression (Lemme *et al.*, 2011; Son *et al.*, 2012; Reck *et al.*, 2015; Hagen; Son, 2017). Transformation efficiency in biofilms is typically less than 0.1% (Y. Li *et al.*, 2001), while even under very favorable conditions no more than 10-50% of cells naturally express *comX* (Lemme *et al.*, 2011; Son *et al.*, 2012). In addition, the expression of *comX* can be bimodal or unimodal in the population, depending on the exogenous signals present, the growth phase and the environment (Son *et al.*, 2012; Shields; Burne, 2016). Post-translational regulation of ComX also appears to generate heterogeneity, as high levels of *comX* mRNA do not assure robust activation of *comY* (Seaton *et al.*, 2011). As with many other bacterial regulatory proteins (Inobe; Matouschek, 2008), ComX levels in *S. mutans* are modulated post-translationally by an ATP-dependent protease system composed of MecA and ClpCP (Tian *et al.*, 2013; Dong *et al.*, 2014; Dufour *et al.*, 2016). The MecA/ClpCP complex inhibits competence by targeting and degrading ComX, as it does in streptococci of the salivarius, mitis and pyogenic groups (Biornstad; Havarstein, 2011; Boutry *et al.*, 2012; Wahl *et al.*, 2014; Y. H. Li; Tian, 2017). However, the function of MecA/ClpCP within the *S. mutans* competence pathway, and particularly its role in cell-to-cell heterogeneity and the bimodal and unimodal competence behaviors, has not been explored in detail.

Figure 1 summarizes the competence regulatory pathway in *S. mutans* (Smith; Spatafora, 2012; Tian *et al.*, 2013; Shanker; Federle, 2016). ComX activates late competence genes that include the nine-gene operon *comYA-I*, which contains seven genes that are required for transformation (Merritt *et al.*, 2005).Transcription of *comX* can be triggered by either of two quorum sensing peptides: CSP (competence stimulating peptide) or XIP (SigX-inducing peptide). The efficacy of these peptides is sensitive to environmental factors, including pH, oxidative stress, carbohydrate source, and the peptide content of the medium.

**Figure 1.**
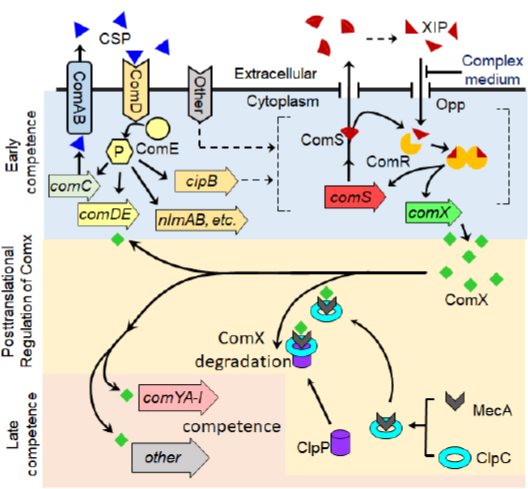
*Streptococcus mutans* regulates genetic competence through multiple layers of control (Smith; Spatafora, 2012; Tian et al., 2013; Shanker; Federle, 2016). Two quorum signals (CSP and XIP), together with other environmental inputs, drive the master competence regulator ComX, which is post-translationally regulated by the MecA/ClpCP proteolytic system. The peptide CSP (competence stimulating peptide) is detected by the ComDE two component system, leading to phosphorylation of the response regulator ComE, which activates transcription of bacteriocin genes such as *cipB*. Through a pathway not yet understood, activation of *cipB* is integrated with other environmental cues to stimulate the ComRS system, which is the immediate regulator of *comX*. ComX, also called SigX, is an alternative sigma factor that directly controls the nine-gene operon *comYA-I* and other genes required for transformation. The ComRS system includes the peptide ComS and the cytosolic receptor ComR. ComS is the precursor for the quorum-sensing peptide XIP (SigX-inducing peptide). In defined growth medium (lacking assorted small peptides), extracellular XIP is imported by the Ami/Opp permease and interacts with ComR to form a transcriptional activator for both *comS* and *comX.* As a result *comX* is uniformly (population-wide) activated in defined media containing XIP. In complex media, which are rich in assorted small peptides, extracellular XIP does not activate *comS* and *comX;* in this case XIP (or possibly its precursor ComS) is proposed to interact with ComR intracellularly, so that both *comS* and *comX* are driven by the bistable, intracellular transcriptional feedback loop involving *comS* and the ComRS complex (Son et al., 2012). As a result *comX* is heterogeneously (bimodal in population) activated in complex media. The MecA/ClpCP system provides posttranslational regulation of ComX: The adapter protein MecA interacts with ClpC to target ComX for degradation by the protease ClpP.

CSP is derived from the ComC precursor, processed to a final length of 18 aa and exported to the extracellular medium. *S. mutans* detects CSP through the ComDE two-component signal transduction system (TCS), which directly activates multiple genes involved in bacteriocin biogenesis, secretion and immunity. However, *S. mutans* ComDE does not directly activate *comX*. Instead, the ComRS system is the immediate regulator of *comX* in the mutans, salivarius, bovis, and pyogenes groups of streptococci (Mashburn-Warren *et al.*, 2010). The ComRS system consists of the cytosolic receptor ComR and the 17-aa peptide ComS, which is processed by an unknown mechanism to form the 7-aa XIP. Extracellular XIP is imported by the oligopeptide permease Opp and interacts with ComR to form a complex that activates the transcription of *comS* and *comX*. Exogenous XIP induces *comX* efficiently in chemically defined media lacking small peptides (such as FMC or CDM (Mashburn-Warren *et al.*, 2010; Son *et al.*, 2012)), leading to population-wide induction of *comX* at saturating XIP levels. However, XIP elicits no induction of *comX* in complex growth media containing small peptides, possibly owing to peptide competition with XIP for uptake by Opp. Interestingly, the CSP peptide signal has a different action than XIP, as it activates *S. mutans comX* only in complex growth media containing small peptides. It elicits no activity from *comX* in defined media that lacks small peptides, even though CSP stimulates the ComDE TCS (leading to bacteriocin production) under these conditions. In addition, the *comX* response to CSP is bimodal in the population, with no more than 50% of cells expressing *comX* at saturating CSP concentrations (Son *et al.*, 2012).

Consequently, the activation of *comX* in a population of *S. mutans* can exhibit two types of heterogeneity: a unimodal distribution when stimulated by exogenous XIP and a bimodal distribution when stimulated by exogenous CSP. Only the bimodal behavior requires an intact *comS*, whereas only the unimodal behavior requires the oligopeptide permease *opp.* We previously posited that these different behaviors are two modes of operation of the transcriptional feedback loop associated with *comS*, which encodes its own inducing signal. In the unimodal case the cells import and respond to exogenous XIP, whereas in the bimodal case XIP import is blocked, leaving each cell to respond to its intracellular ComS (or XIP). The first mode allows a generally uniform, population-wide activation of *comX*, but the second mode leads to noisy, positive feedback dynamics in both *comS* and *comX* (Son *et al.*, 2012; Hagen; Son, 2017).

The mechanism of posttranslational control of ComX by MecA/ClpCP in *S. mutans* resembles that in pyogenic and salivarius streptococci, to which *S. mutans* MecA is closely homologous (Boutry *et al.*, 2012; Wahl *et al.*, 2014). *S. mutans* MecA is a 240 aa adapter protein that interacts with ComX and ClpC to form a ternary complex that sequesters ComX and targets it for ATP-dependent degradation by the ClpP protease (Tian *et al.*, 2013; Dong *et al.*, 2014). MecA/ClpCP similarly controls the master competence regulators ComW in *S. pneumoniae* (Wahl *et al.*, 2014) and ComK in *Bacillus subtilis* (Turgay *et al.*, 1998). In *B. subtilis* MecA was shown to facilitate the ATP-dependent formation of the ClpCP proteolytic complex, which unfolds and degrades both MecA and its ComK target, and then itself dissociates (Mei *et al.*, 2009; Liu *et al.*, 2013). Therefore MecA/ClpCP operates dynamically by continuously turning over MecA as well as its regulatory target if present.

Several studies in *S. mutans* have established that MecA/ClpCP suppresses the activation of *comY* under CSP stimulation, in complex media (Tian *et al.*, 2013; Dong *et al.*, 2014; Dufour *et al.*, 2016). Deletion of *mecA*, *clpC*, or *clpP* increased ComX levels and transformability during growth in complex media and also prolonged the competent state. These studies imply that MecA/ClpCP serves either to suppress *S. mutans* competence or to switch it off as growth progresses, in complex media. Some studies have found the puzzling result that deletion of *mecA* or *clpCP* caused a weaker increase in ComX levels or transformability - or even had no effect at all – in chemically defined media (with added XIP) than in complex media (with CSP) (Boutry *et al.*, 2012; Tian *et al.*, 2013; Dong *et al.*, 2014; Dufour *et al.*, 2016). A subsequent study found that MecA deletion improved *S. salivarius* transformability in defined media, although the difference was attenuated at high levels of XIP stimulation (Wahl *et al.*, 2014).

The possible significance of growth media and the presence of heterogeneity raise the question of how MecA/ClpCP functions within the full competence pathway, in which the XIP and CSP signaling pathways activate *comX* in defined and complex media respectively. Although it seems clear that MecA/ClpCP inhibits *comY* expression by sequestering and degrading ComX, a clearer model of how this regulation integrates with the known *comX* activation pathway, and how it may be overcome when competence is permitted, is still needed. Additional cell density signals (Dufour *et al.*, 2016), as well as XIP-dependent feedback or additional gene products (Wahl *et al.*, 2014), have been proposed as mechanisms for modulating ComX levels via MecA/ClpCP. We have used a single-cell, microfluidic approach to clarify some of these questions and to develop an explicit model of how MecA/ClpCP interacts with the noisy and bimodal mechanisms controlling *S. mutans comX*. Our data lead to a simple quantitative model that reproduces both the population average behavior and the cell-to-cell heterogeneity in *comY* activation.

## Results

### MecA/ClpCP affects transformation efficiency of S. mutans induced by XIP

Supporting Figure S4 shows how deletions in the MecA/ClpCP system affect transformation efficiency of *S. mutans* UA159. Transformability was measured in cells cultured in defined medium (FMC) containing various concentrations of XIP, as indicated. At the highest XIP concentration (1 μM), the transformation efficiencies of the *mecA* and *clpC* deletion mutants were similar to the wild type. This finding is consistent with previous reports for *S. mutans* (Tian *et al.*, 2013; Dong *et al.*, 2014) and *S. thermophilus* (Boutry *et al.*, 2012), where little or no effect of *mecA* deletion on transformability in response to XIP was observed. However the behavior of the mutants diverged at lower XIP concentrations, where deficiency of MecA or ClpCP enhanced transformability. An effect of XIP concentration on the behavior of deletion mutants was also reported for *S. salivarius* (Wahl *et al.*, 2014). Both Δ*clpP* and Δ*clpC* had higher transformation efficiency than the wild type at 10 nM XIP. Surprisingly the *mecA* deletion showed lower transformability than the wild type strain at 100 nM XIP; we note however this strain grows poorly and displays defects in cell morphology and decreased viability. Overall these data confirm that the MecA/ClpCP system interacts with XIP induction of transformability in defined medium. To obtain more detailed insight into XIP stimulation, MecA/ClpCP and *comY* activation, we turned to individual cell studies.

### Activation of comX leads to heterogeneous induction of comY

We used dual fluorescent reporters (P*comX-gfp*, P*comY-rfp*) to compare the activation of P*comX* and P*comY* in individual *S. mutans* supplied with exogenous XIP. Figure 2A shows *S. mutans* UA159 growing in microfluidic channels under a constant flow of defined medium (FMC) that contains 0-2 µM XIP. P*comX* is activated in all cells if the XIP concentration exceeds about 100 nM, and its activation saturates as XIP exceeds about 800 nM. However, very few cells activate P*comY* at XIP concentrations of 400 nM or less, and cells that do activate P*comY* vary widely in their red fluorescence intensity. Even at 1-2 μM XIP, many cells exhibit little P*comY* activity.

**Figure 2.**
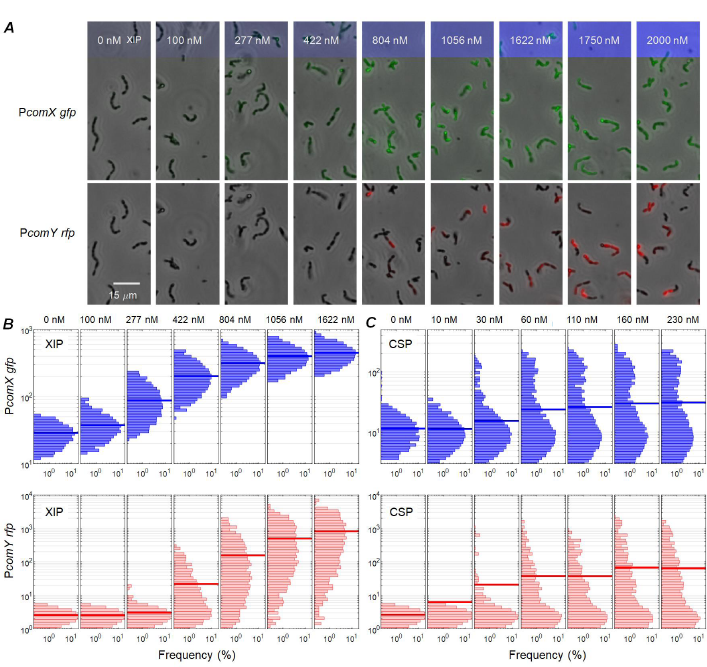
(A) Microscopy images of dual reporter (P*comX-gfp* and P*comY-rfp*) UA159 *S. mutans* growing in microfluidic channels. Cells were supplied with the indicated concentrations of synthetic XIP by a continuous flow of defined medium. Phase contrast images (grayscale) are overlaid with fluorescence images showing P*comX* (GFP, upper) and P*comY* (RFP, lower) activity. (B)-(C) Histograms of *comX* (upper row) and *comY* (lower row) expression in dual reporter UA159 *S. mutans* under two different modes of stimulation. Cells growing in microfluidic channels were supplied with a continuous flow of (B) defined medium containing XIP, or (C) complex medium containing CSP, and were imaged in red (for P*comY*) and green (for P*comX*) fluorescence. XIP and CSP concentrations are indicated along the top of the figures. The length of each horizontal histogram bar indicates the percentage of cells that express at the level indicated. All axes are logarithmic. The thick blue or red bar in each histogram shows the population mean.

Figures 2B-2C show the statistical distribution of P*comX* (GFP, upper rows) and P*comY* (RFP, lower rows) reporter fluorescence for cells in response to exogenous XIP or CSP. Reporter fluorescence was imaged while cells grew in microfluidic channels under continuous flow of defined medium for XIP (Figure 2B), or of complex medium for CSP (Figure 2C). As previously reported (Son *et al.*, 2012), XIP in defined medium elicits a noisy but generally unimodal (population-wide) *comX* response. In contrast, CSP in complex medium elicits a much noisier, bimodal (double peaked distribution) *comX* response. For both CSP and XIP stimulation, the response of P*comY* is highly heterogeneous. Even the highest concentrations of CSP and XIP, which saturate the response of P*comX*, incompletely activate P*comY* in the population; the P*comY* expression levels in individual cells span 2-3 orders of magnitude above the baseline. These data suggest that post-translational regulation of ComX increases cell-to-cell heterogeneity in *comY* expression, which adds to the noise in the *comX* response to CSP or XIP stimulation.

### The MecA/ClpCP system inhibits the comY response to XIP and increases its noise

To test whether the MecA/ClpCP proteolytic system affects ComX function in defined medium, and to assess its effect on noise in *comY* expression, we compared *comX* and *comY* expression in dual reporter strains in the wild-type (UA159) and Δ*mecA* genetic backgrounds. Figure 3 shows P*comY* activity in individual cells that were stimulated by XIP in planktonic culture in defined medium and then imaged on glass slides. Similar results were obtained for cells growing in microfluidic flow channels. Deletion of *mecA* altered the P*comY* response in two ways. First, the Δ*mecA* strain responded more strongly to XIP than did the wild type. Unlike the wild-type genetic background, the Δ*mecA* cells showed high median P*comY* expression, exceeding the baseline level at XIP concentrations greater than about 200 nM. Second, deletion of *mecA* reduced noise in *comY* expression (Figure 3C, 3D). Although *comY* and *comX* expression correlated positively in UA159, the correlation was partially obscured by the noisy behavior of *comY*. In contrast, *comY* expression increased systematically as *comX* expression increased in the Δ*mecA* strain. Despite some noise in *comY*, a roughly proportional relationship can be discerned in the data of Figure 3D, but not in Figure 3B. (The upward curvature in Figure 3D results from the logarithmic horizontal axis.) The nearly linear correlation between *comY* and *comX* in the *ΔmecA* mutant suggests that, in the absence of MecA/ClpCP, ComX activates *comY* in a direct and predictive fashion.

**Figure 3.**
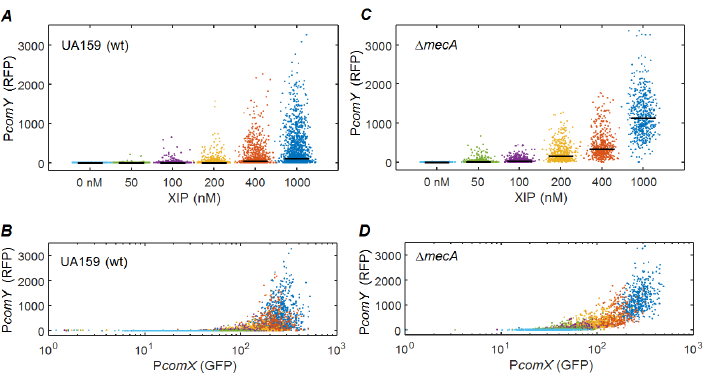
Comparison of noise in *comY* activation by XIP in (A)-(B) UA159 (wild type) and (C)-(D) Δ*mecA* deletion strain of *S. mutans*. Each point shows the RFP fluorescence of one cell that was incubated with XIP at the indicated concentration and then imaged on a glass slide. (A) and (C) show the dependence of *comY* expression on XIP stimulus in the two strains. The horizontal bar indicates the median expression. (B) and (D) show the correlation between *comY* and *comX* expression within individual cells, with the point colors indicating the XIP concentration in the same color code as (A) and (C).

The effect of MecA on noise in *comY* is also seen in histograms of *comY* expression at given *comX* expression levels. Supporting Figure S2 shows *comY* histograms for cells growing in microfluidic channels with flowing defined medium and XIP, binned according to their *comX* activity. Both at high and low *comX* activity, the shape of the *comY* histograms is qualitatively different in the two strains. The deletion of *mecA* qualitatively alters the relationship between *comY* and *comX* expression in defined medium with addition of XIP.

### comY and comX expression are simply correlated in the absence of MecA

As is common for bacterial protein expression (Taniguchi *et al.*, 2010), the histograms of P*comY* expression (Supporting Figure S2) resemble a gamma distribution Γ(*n* | *A,B*), a two-parameter continuous probability distribution that can be interpreted in terms of sequential, stochastic processes of transcription and translation (see *Methods*). This finding, together with the roughly linear correlation between *comY* and *comX* activity in the Δ*mecA* strain (Figure 3D), motivates a simple mathematical model for *comX/comY* in the absence of MecA/ClpCP. The model is described in the *Methods*: *comY* is activated in a mostly linear (but saturating) fashion by *comX* on average, but is also subject to stochasticity. The *comY* activity in a given cell is thus a random variable drawn from a gamma distribution whose parameters are determined by the P*comX* activity in the cell. The model has four parameters, which we obtained through a maximum likelihood fit to the Δ*mecA* individual cell RFP and GFP fluorescence data of Figure 3D. We then used these parameters to generate a stochastic simulation of the model for comparison to the data.

Figure 4 compares the Δ*mecA* experimental data (Figure 4A, 4B) with a simulation of the model (Figure 4C, 4D). The model accurately reproduces both the population-averaged *comY* response and its cell-to-cell variability. This result indicates that in the absence of MecA/ClpCP regulation of ComX, *comY* can be modeled as a typical noisy gene whose average activation is proportional to the concentration of active ComX protein.

**Figure 4.**
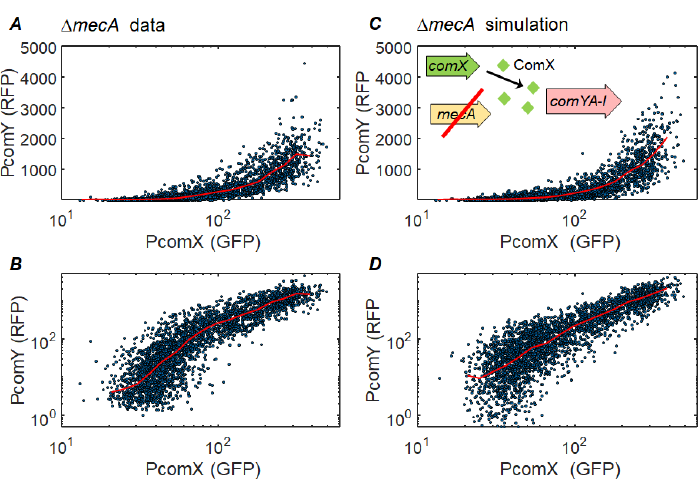
Relation between P*comY* and P*comX* activity in the Δ*mecA* deletion strain, in response to XIP: (A)-(B) Experimental data of Figure 3, showing correlation between P*comX* and P*comY* expression in Δ*mecA* cells subject to a range of XIP concentrations; (C)-(D) simulation of a stochastic model for *comX* activation of *comY*. The stochastic model, described in *Methods*, assumes that P*comX* activity within each cell directly determines the probability distribution for P*comY* activation in that cell. The upper and lower rows show the same data on linear and logarithmic vertical scales, respectively. The red curves show the median response. Supporting Figure S2 shows an additional comparison between the data and an alternative model in which the environmental XIP concentration, rather than P*comX* activity, determines the probability distribution for P*comY* activation. Model parameters are given in the legend to Supporting Figure S2.

A plausible alternative model is that extracellular XIP concentration, rather than P*comX* activity *per se*, controls *comY* expression in Δ*mecA*. The simulation shown in Supporting Figure S3 indicates that the best fit of this model significantly overestimates the noise in P*comY*. In short, modeling suggests that the P*comX* activity of a Δ*mecA* cell is a straightforward predictor of its P*comY* activity, and is also a better predictor than is the XIP concentration.

### Different deletions in MecA/ClpCP produce different noise and threshold behaviors in comY

To determine which elements of the MecA/ClpCP system affect sensitivity and noise in *comY*, we measured P*comY* and P*comX* activity in the UA159, Δ*mecA*, Δ*clpC* and Δ*clpP* genetic backgrounds (Figure 5). All strains carried the dual fluorescent reporters and were imaged in microfluidic chambers while supplied with a continuous flow of defined medium containing XIP. In all strains, P*comY* was more strongly activated at higher XIP concentrations where Pc*omX* expression was higher, although noise and sensitivity varied among the different strains (Figure 5A). All strains showed a similar dependence of P*comX* activity (GFP) on XIP concentration (Figure 5B). In the relation between *comY* and *comX* expression, the UA159 (wild type) showed a more pronounced threshold in the onset of *comY* activation, at a *comX* level near 300 units, and much greater noise in *comY* expression. The *clpP* deletion strain, in which the MecA/ClpC complex can presumably bind, but not degrade, ComX, showed slightly less noisy *comY* expression than the wild type and *comY* was somewhat more readily activated. Deletion of *clpC*, or especially *mecA*, reduced *comY* noise significantly, such that the population was almost uniformly activated when P*comX* expression was strong, near 1 μM XIP. Therefore, the interaction between MecA/ClpC and ComX, as well as the proteolytic action of ClpP on that complex, contribute to noise in *comY* expression and also suppress the ability of *comX* expression to elicit the *comY* response. Similar data were obtained when cells were grown in static medium and image while dispersed on glass slides.

**Figure 5.**
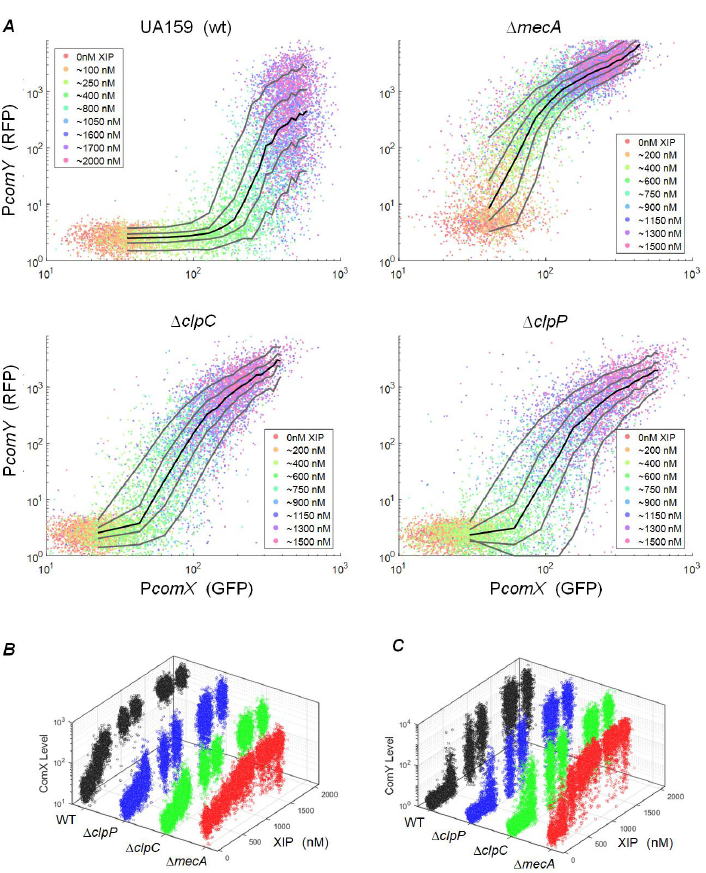
(A) Effect of *mecA/clpCP* deletions on the correlation between *comY* and *comX* activation. For each of the four strains (UA159, Δ*mecA*, Δ*clpP*, Δ*clpC*) each point shows the P*comY* and P*comX* activity of one cell, as measured by RFP and GFP reporters respectively. Cells were imaged while growing in microfluidic channels that were supplied with a continuous flow of defined medium that contained XIP concentrations as indicated by the point color. Approximately 1000 cells of each strain were imaged at each XIP concentration. Solid lines indicate the 10, 30, 50, 70, and 90^th^ percentiles of P*comY* activity. Cell autofluorescence contributes a background red fluorescence that is typically 1-5 fluorescence units in most experiments. Cell autofluorescence contributes a green background that is typically 20-30 fluorescence units. Supporting Figure S6 shows the same data on linear axes. (B)-(C) Scatterplots showing individual cell *comX* and *comY* expression versus exogenously added XIP in the four strains.

### The role of MecA alone can be modeled by simple sequestration of ComX

A detailed model for the regulation of ComX by MecA/ClpCP must include the formation of the MecA/ClpC/ComX ternary complex, as well as the kinetics of ComX and MecA degradation by ClpP. Both of these mechanisms are absent in the Δ*clpC* strain, although the binary interaction of MecA with ComX is present. Therefore, we tested whether a binary sequestration (MecA + ComX) model could reproduce our data for the activation of *comY* by ComX in the *ΔclpC* strain. In this model, described in *Methods*, individual ComX molecules are presumed to be tightly sequestered by individual MecA molecules, leaving them unavailable to stimulate *comY* transcription. Then the probability distribution for the *comY* expression of a cell becomes determined not by its *comX* activity alone, but by the excess of ComX over MecA copy numbers. We modeled the MecA copy number as a random variable drawn from a gamma probability distribution; the activation of *comY* by the available (unsequestered) ComX is modeled as in Figure 4. The MecA probability distribution is presumed to be independent of XIP, consistent with our mRNA measurements showing no effect of XIP on *mecA*, *clpC* or *clpP* expression (Supporting Table ST1). Fitting this MecA model to the Δ*clpC* data therefore requires only a two-parameter fit for the gamma distribution parameters, which we obtained by maximum likelihood comparison of the data and model.

Figure 6 compares the Δ*clpC* data with a stochastic simulation of this model. The *comY - comX* correlation closely resembles the experimental data, both in its average trend and its noise. These results show that the higher *comY* expression noise that is observed in the Δ*clpC* strain, compared to the Δ*mecA* strain, is consistent with a mechanism where MecA suppresses *comY* response by sequestering ComX. Fitting the model to the data provides the probability distribution of the MecA copy number, Figure 6C, where MecA is measured in units of equivalent P*comX* activity. Cell-to-cell variability in MecA copy number is then a source of variability in *comY* expression.

**Figure 6.**
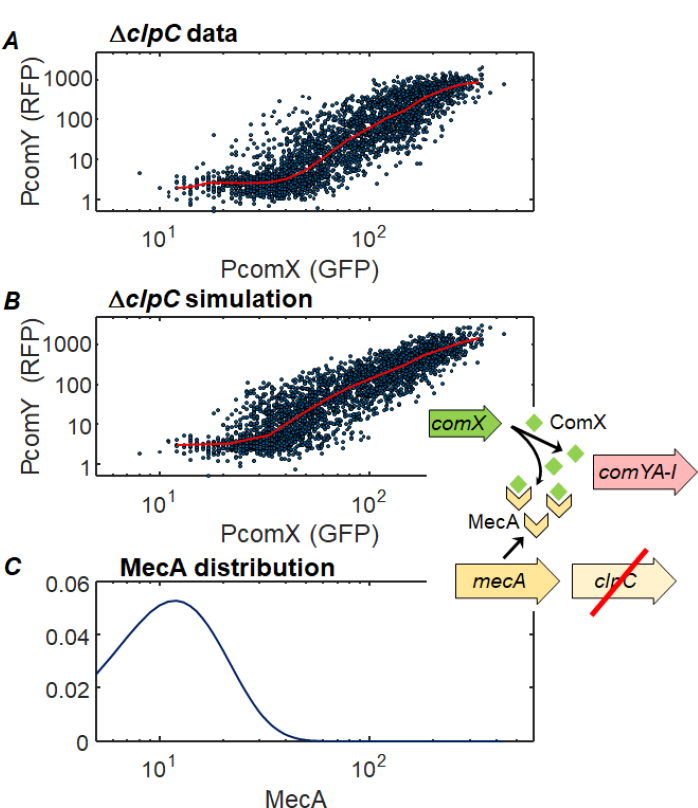
Data and model for MecA sequestration of ComX. (A) Experimental measurement of *comX* activation of *comY* in dual-reporter (P*comX-gfp*, P*comY-rfp*) Δ*clpC* cells. Cells were provided 50-1000 nM XIP (defined medium) and then imaged on glass slides. The solid red line is the median P*comY* response. (B) Simulation of a stochastic model (see *Methods*) in which the activation of P*comY* in each cell is determined by the excess of the cell’s P*comX* activity over its MecA level, where MecA levels obey a gamma statistical distribution. The simulation in (B) uses the MecA distribution (parameters A = 3.61, B = 4.53) that maximizes the likelihood of the data in (A). The baseline or background in *comY* is modeled by gaussian noise of 3 ± 0.8 red fluorescence units. (C) The statistical distribution of MecA levels used in generating the simulation of (B). MecA levels are referenced to P*comX* expression levels: A MecA copy number of 10 implies MecA exactly sufficient to sequester all of the ComX produced when PcomX-gfp expression is at the level 10.

### CSP and XIP stimulation produce similar correlations between comX and comY activation

Previous studies have demonstrated that deletions of *mecA* or *clpCP* enhance *comY* expression upon stimulation with CSP in complex media (Tian *et al.*, 2013; Dong *et al.*, 2014). Our data show with single cell resolution that the same deletions also affect the response to XIP in defined media. These findings raise the question of whether, in the presence of MecA/ClpCP, the activation of *comY* by ComX may be similar regardless of how *comX* transcription is induced, whether by XIP or CSP. Figure 7 compares single cell measurements of *comX* and *comY* activity with CSP and XIP respectively. Precise quantitative comparison of the two response curves is complicated by the stronger green auto-fluorescence of cells in complex medium, which shifts the horizontal axis of the CSP data. Further, CSP appears to induce a slightly noisier *comY* response than does XIP, possibly in connection with feedback behavior in the ComDE system (Son *et al.*, 2015a). However the data verify a generally similar behavior in both conditions: *comY* responds in threshold fashion to activation of *comX*, and *comY* activation is highly heterogeneous in the population, even among cells with the highest *comX* activity.

**Figure 7.**
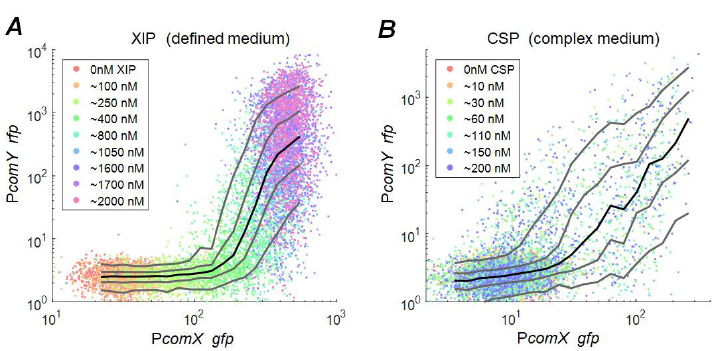
Comparison of *comX*/*comY* correlation in UA159 background in response to XIP and CSP. Cells were imaged while adhered in microfluidic flow channels supplied with continuous flow of (A) defined medium containing XIP or (B) complex medium containing CSP. Solid lines indicate the 10, 30, 50, 70, and 90^th^ percentiles of P*comY* activity. Horizontal scales are not strictly equivalent owing to higher autofluorescence baseline of cells in the complex media.

## Discussion

The MecA/ClpCP proteolytic system is well conserved as a negative regulator of genetic competence across streptococcal groups and in other naturally competent species, including *B. subtilis* (Liu *et al.*, 2013). However, while mechanistic studies of MecA/ClpCP have provided a clear description of its action, they have not fully resolved the question of how MecA/ClpCP contributes to competence regulation. Several authors have proposed that MecA/ClpCP serves either to suppress or terminate the competent state. For *B. subtilis*, Turgay *et al.* proposed that MecA/ClpCP degradation of the ComK competence regulator provides a ‘timing’ function by limiting synthesis of the auto-activating ComK regulator, thus permitting escape from the competent state (Turgay *et al.*, 1998). Dufour *et al*. proposed a similar model for *S. mutans*, in which the sequestration and degradation of free ComX by MecA/ClpCP forces an exit from the competent state late in growth, when the transcription of *comX* is repressed (Dufour *et al.*, 2016). Wahl *et al*. proposed that *S. salivarius* MecA/ClpCP serves a ‘locking’ function, preventing the cell from entering the competent state under inappropriate conditions, such as early in the growth phase (Wahl *et al.*, 2014). Wahl *et al*. argued that at low XIP concentrations proteolytic degradation of ComX prevents competence, but that high XIP concentrations may alleviate this repression, possibly by overwhelming the proteolytic capacity or by activating another, unidentified gene product.

Both the ‘locking’ and ‘timing’ models interpret MecA/ClpCP as a mechanism for suppressing activation of *comY* when *comX* expression is weak. Our data are consistent with this description. Moreover, our data show that this suppression can be described by the simplest model in which an intracellular pool of MecA intercepts available ComX, sequestering it and blocking its otherwise straightforward activation of *comY*. Such a model quantitatively fits the data on the *clpC* mutant, in which MecA can sequester ComX but *clpP* proteolysis is absent. If the MecA copy number obeys a gamma probability distribution, as is common for bacterial proteins, then the model reproduces both the average relationship between *comY* and *comX* expression and the cell-to-cell variability in that expression. Therefore, the response of *comY* in individual *clpC* and *mecA* cells can be understood solely in terms of the P*comX* activity and MecA copy number distribution. The behavior of the late competence genes in these mutants can be understood without positing any role for XIP other than as a stimulus for P*comX*.

In addition, our single cell data show that the MecA/ClpCP system substantially enhances the noise (cell-to-cell heterogeneity) in *comY* expression when *comX* is activated. Even at high XIP concentrations that saturate *comX* expression, *comY* expression levels within the UA159 population span a range extending three orders of magnitude above the baseline; by contrast, the deletion mutants all express *comY* with far less noise at high XIP concentrations. Our modeling indicates that cell-to-cell variability in the MecA copy number in wild type cells, together with the proteolytic action of ClpP (which reduces MecA and ComX copy numbers) adds to noise that is generated upstream by the pathways that activate *comX*. The resulting noisy threshold effect is very similar to the toxin/antitoxin competition that generates phenotypic heterogeneity in bacterial persistence (Rotem *et al.*, 2010), or to a sequestration-induced threshold model for non-linear gene regulation (Buchler; Cross, 2009).

A clear understanding of the role of MecA/ClpCP has perhaps been complicated by early reports that deletion of *mecA* or *clpC* increased transformability or ComX protein levels under CSP stimulation (in complex medium), but not under XIP stimulation (in defined medium). Our data confirm in detail that the MecA/ClpCP system affects signaling from *comX* to *comY* in defined medium. In fact, as the sequestration model described above is indifferent to whether *comX* is stimulated by exogenous XIP or CSP, we expect that signaling from *comX* to *comY* should be similar in both CSP/complex medium and in XIP/defined medium. Figure 7 suggests that the relationship is very similar.

This finding suggests that the MecA/ClpCP system acts continuously to suppress ComX levels, regardless of the extracellular inputs driving *comX* expression. A model where MecA/ClpCP performs this task in relatively steady fashion is consistent with findings that *S. mutans* MecA and ClpCP protein levels did not differ in complex and defined medium (Dong *et al.*, 2014), that MecA induction showed little change during *S. suis* competence (Zaccaria *et al.*, 2016), and that *S. mutans mecA/clpCP* mRNA levels are insensitive to XIP inputs (Supporting Table ST1). Thus competence will be suppressed when *comX* is weakly expressed due to insufficient CSP or XIP early in growth (‘locking’ behavior). Competence will also be suppressed when *comX* is weakly expressed late in growth due to inefficient CSP/XIP signaling. Falling extracellular pH late in the growth phase suppresses competence signaling by CSP and XIP (Guo *et al.*, 2014; Son *et al.*, 2015b), which may allow MecA/ClpCP to shut down the competent state (‘timing behavior’).

Consequently the sequestration mechanism can provide both ‘timing’ and ‘locking’ functions. The simulations in Figure 4 and Figure 6 are based on simple equilibrium models that address only the effects of sequestration by MecA on the pool of free ComX, omitting the kinetic effects of ClpP unfolding and degradation of ComX and MecA. A model that includes ClpP proteolysis is much more complicated, as it must include the sequential binding steps that are associated with the formation of the ternary complex, binding of ClpP, and the breakdown of both MecA and ComX. The binding and kinetic parameters of the model cannot be determined from our data; however we can construct a reasonably tractable model for the full system by simplifying the complex regulatory mechanism that is outlined in the literature (Mei *et al.*, 2009). Supporting Figure S7 describes a simplified kinetic model that can rationalize some of the observations in our data, including the finding that deletion of *clpC* or *clpP* did not eliminate the *comX* threshold that is required for *comY* activation, and that only the *mecA* deletion eliminated the threshold and sharply reduced the noise in *comY*. Supporting Figure S7 shows that simulations from such rough models can reproduce key differences in *comX-comY* threshold behavior observed among the mutants studied here.

We note that a MecA copy number distribution that has higher mean but is narrower than that of Figure 6C would still provide the same ‘timing’ or ‘locking’ function without introducing as much noise in *comY*. The evident width of the distribution therefore suggests that the organism may benefit from greater noise. The competence pathway in *S. mutans* is linked to several stress-induced behaviors that are heterogeneous in the population, including competence, lysis and a persister phenotype (Perry *et al.*, 2009; LeungAjdic *et al.*, 2015; Leung *et al.*, 2015). A link between quorum controlled behavior and phenotypic heterogeneity has often been noted in bacterial gene regulation. In other organisms, such as *B. subtilis*, complex pathways that integrate intracellular and extracellular signaling mechanisms with stochastic gene expression often generate phenotypic heterogeneity, distributing stress response behaviors such as competence and sporulation among different individuals in the population (Grote *et al.*, 2015). Interestingly, propidium iodide staining of individual *S. mutans* indicates that *comX*-driven lysis is decoupled from *comX*-driven competence (Supporting Figure S5). While higher *comX* expression increases the probability of cell lysis, the most highly expressing cells (which are more likely to express *comY*) actually show less evidence of lysis. Accordingly, the MecA/ClpCP system may provide a bet-hedging advantage to an *S. mutans* population by providing an additional, stochastic switching point in the regulatory pathway from stress conditions to transformability.

Our data show that the action of the *S. mutans* MecA/ClpCP system can be quantitatively understood, at the level of individual cell behavior, within a very simple threshold mechanism. As the MecA/ClpCP system is widely conserved this finding raises the question of whether MecA/ClpCP also generates a heterogeneity advantage in other organisms such as *S. pneumoniae*, in which competence regulation is more straightforward and the *comX* bimodality mechanism is absent. Our data also highlight the long standing question of whether by combining cooperative behaviors of quorum signaling with deliberately noisy intracellular phenomena such as MecA and ComRS, *S. mutans* can achieve some form of optimum balance between socially-driven, environmentally-driven and purely stochastic behavior in competence regulation.

## Experimental Procedures

### Preparation of reporter strains

*S. mutans* strains and deletion mutants (Table 1) harboring green fluorescent protein (*gfp*) and/or red fluorescent protein (*rfp*) reporter genes fused to the promoter regions of *comX* (P*comX-gfp*) and *comY* (P*comYA-rfp*) were grown in brain heart infusion medium (BHI; Difco) at 37°C in a 5% CO_2_, aerobic atmosphere with either spectinomycin (1 mg mL^−1^), erythromycin (10 µg mL^−1^), or kanamycin (1 mg mL^−1^). P*comX-gfp* was directly integrated into the chromosome of *S. mutans* (denoted XG) by amplifying a 0.2-kbp region comprising P*comX* using primers that incorporated *Xba*I and *Spe*I sites (Table 2). This was fused to a *gfp* gene that had been amplified with primers engineered to contain *Spe*I and *Xba*I sites from the plasmid pCM11 (Lauderdale *et al.*, 2010; Son *et al.*, 2012), and inserted into the *Xba*I site on pBGE (Zeng; Burne, 2009). P*comYA-rfp* was constructed in shuttle vector pDL278 (LeBlanc *et al.*, 1992) by amplification of a 0.2-kbp region containing P*comY* with *Hin*dIII and *Spe*I site-containing primers and fusing with the *rfp* gene reporter fragment amplified from plasmid pRFP (Bose *et al.*, 2013), using primers that incorporated *Spe*I and *Eco*RI sites. The ligation mixtures were transformed into competent *S. mutans* (strain designated YR) and into the XG strain (denoted XG&YR). Additionally, to study the role of MecA/ClpCP on P*comY* expression, both the XG integration vector and the YR shuttle vector were transformed into strains harboring non-polar (NPKmR) antibiotic resistance cassette replacements of *mecA* (this study), *clpC* or *clpP* (J. A. C. Lemos; Burne, 2002). Plasmid DNA was isolated from *Escherichia coli* using a QIAGEN (Chatsworth, Calif.) Plasmid Miniprep Kit. Restriction and DNA-modifying enzymes were obtained from Invitrogen (Gaithersburg, Md.) or New England Biolabs (Beverly, Mass.). PCRs were carried out with 100 ng of chromosomal DNA using Taq DNA polymerase. PCR products were purified with the QIAquick kit (QIAGEN). DNA was introduced into *S. mutans* by natural transformation and into *E. coli* by the calcium chloride method (Cosloy; Oishi, 1973).

**Table 1:** Strains used in this study. Em^r^, erythromycin; NPKm^r^, non-polar kanamycin; Sp^r^, spectinomycin.

| *S. mutans* strains | Characteristic(s) | Source |
| --- | --- | --- |
| WT (UA159) | *S. mutans* wild-type strain | ATCC 700610 |
| XG | $P_{comX}$-*gfp* integrated into UA159, Em<sup>r</sup> | This study |
| YR | UA159 harboring $P_{comY}$-*rfp*, Em<sup>r</sup> | This study |
| XG&YR | XG harboring $P_{comY}$-*rfp*, Sp<sup>r</sup> | This study |
| XG&YR&$\Delta mecA$ | $\Delta mecA$::NPKm<sup>r</sup> into XG&YR | This study |
| XG&YR&$\Delta clpC$ | $\Delta clpC$::NPKm<sup>r</sup> into XG&YR | This study |
| XG&YR&$\Delta clpP$ | $\Delta clpP$::NPKm<sup>r</sup> into XG&YR | This study |

**Table 2:**
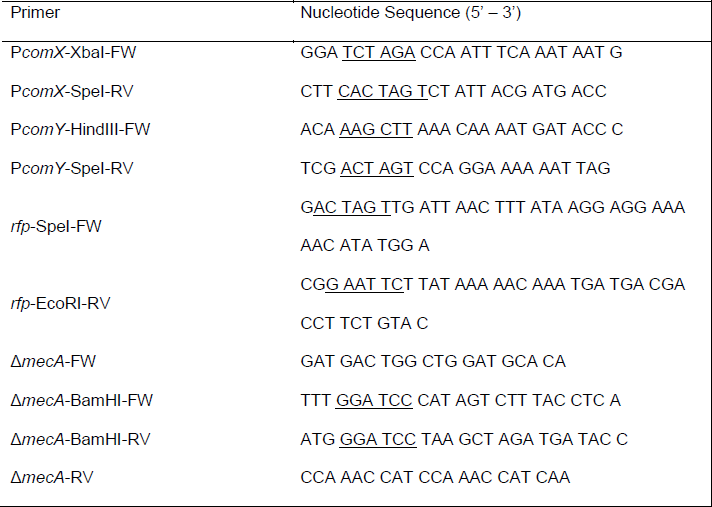
Primers used in this study. Underline of nucleotide sequence denotes respective restriction enzyme site.

### Competence Peptides

Synthetic CSP (sCSP, aa sequence = SGSLSTFFRLFNRSFTQA), corresponding to the mature 18 aa peptide (Hossain; Biswas, 2012) was synthesized by the Interdisciplinary Center for Biotechnology Research (ICBR) facility at the University of Florida and its purity (95%) was confirmed by high performance liquid chromatography (HPLC). sCSP was reconstituted in water to a final concentration of 2 mM and stored in 100 μL aliquots at −20°C. Synthetic XIP (sXIP, aa sequence = GLDWWSL), corresponding to residues 11-17 of ComS, was synthesized and purified to 96% homogeneity by NeoBioSci (Cambridge, MA). The lyophilized sXIP was reconstituted with 99.7% dimethyl sulfoxide (DMSO) to a final concentration of 2 mM and stored in 100 μL aliquots at −20°C.

### Microfluidic mixer design and fabrication

Microfluidic devices were fabricated by the soft lithography method of molding a transparent silicon elastomer (polydimethylsiloxane) on a silicon master (Sia; Whitesides, 2003). The master was made from a silicon wafer through conventional photolithographic processing. Details of the fabrication method and the devices were described previously (Jeon *et al.*, 2000; Son *et al.*, 2012; Son *et al.*, 2015b). Our microfluidic device consisted of nine parallel flow chambers (each 15 µm deep and 400 µm wide), as shown in Supporting Figure S1. Three inlet channels supplied media containing different concentrations of signal peptides, delivered by syringe pumps into the device. The design has a mixing network that generates nine streams containing different admixtures of the three input solutions. These streams flow through the nine cell chambers in which *S. mutans* are adhered to the lower, glass window. The device also has two side channels: one for the control of fluid inside the device and the other for injection of different solutions into the cell chambers. Two-layer lithography allows air-pressure control of these side channels during cell loading and injection of different solutions (Unger *et al.*, 2000).

### Microfluidic experiments

Overnight cultures grown in BHI with antibiotic selection were washed and diluted 20-fold in fresh medium, which was either chemically defined medium (FMC) (Terleckyj *et al.*, 1975; De Furio *et al.*, 2017) or a complex medium that consisted of 1/3 of BHI (BD) and 2/3 of FMC by volume. Cultures were then incubated at 37°C in a 5% CO_2_, aerobic atmosphere. When OD_600_ reached 0.1 - 0.2, cells were sonicated at 30% amplitude for 10 sec (Fisher FB120) to separate cell chains and then loaded into the microfluidic device. Each cell chamber was continuously perfused with fresh medium containing different amounts of synthetic XIP (0 - 2 µM) or synthetic CSP (0 - 1 µM). The XIP or CSP concentration in each flow channel was generated by the mixture of three different inlet media in the mixing network in the device. A trace amount (0 - 10 ng/mL) of far-red fluorescent dye (Alexa Fluor 647) was added to each of the three inlet media in proportion to its signal molecule concentration, so that the concentration of signal molecule in each chamber could be calculated. After 2.5 h of incubation time, fresh medium containing 100 μg mL^−1^ of rifampicin was flowed through all cell chambers to halt GFP and RFP translation. Cell chambers were then incubated an additional 3 h to allow the full maturation of RFP. Cells were imaged in phase contrast and in green and red fluorescence using an inverted microscope (Nikon TE2000U) equipped with a computer controlled motorized stage and cooled CCD camera.

### Single cell image analysis

Custom Matlab software was used to analyze the expression of the *gfp* and *rfp* reporters in individual cells from overlaid phase contrast, GFP, and RFP images (Kwak *et al.*, 2012). The software first segments individual cells from the cell chain based on the phase contrast image, then finds the concentration of GFP and RFP by correlating the intensity of the phase contrast image with its GFP and RFP fluorescence intensity. This gives a unitless parameter (denoted *R*) that is proportional to the intracellular concentration of GFP or RFP. The GFP or RFP expression levels shown in the data figures are the *R-*values for green or red cell fluorescence respectively.

### Transformation efficiency

Overnight cultures of selected strains were diluted 1:20 into 200 µL of FMC medium in polystyrene microtiter plates. Cells were grown to OD_600_ = 0.15 in a 5% CO_2_ atmosphere. When desired, 300, 500 or 1000 nM of sXIP was added and cells were incubated for 10 min. Then 0.5 µg of purified plasmid pIB184, which harbors an erythromycin resistance (Erm^R^) gene, was added to the culture. Following 2.5 h incubation at 37°C, transformants and total CFU were enumerated by plating appropriate dilutions on BHI agar plates with and without the addition of 1 mg mL^−1^ erythromycin, respectively. CFU were counted after 48 h of incubation. Transformation efficiency was expressed as the percentage of transformants among the total viable cells. The data presented are averages of two independent experiments that each included three biological replicates.

### mRNA levels for mecA, clpCP, and com genes

Data for the analysis of relative mRNA levels for *mecA*, *clpCP* and *com* genes was taken from RNA-Seq analysis completed on strain UA159 (Kaspar *et al* 2018, in preparation). The wild-type strain was grown in FMC medium to OD_600_ = 0.2, at which time either a final concentration of 2 M XIP or vehicle control (1% DMSO) was added. The strains were then allowed to grow to mid-exponential phase (OD_600_ = 0.5) before harvesting. From the analyzed RNA-Seq data, total read counts for each selected gene were found from three biological replicates and RPKM (reads per kilobase per million) calculated under each condition. Finally, ratios for mRNA levels were found by using the normalized RPKM data and by setting *mecA* levels to 1.0. The data files used in this study are available from NCBI-GEO (Gene Expression Omnibus) under accession no. GSE110167.

### Stochastic model for MecA regulation of comX

We used the gamma statistical distribution to model cell-to-cell variability (noise) in the activation of *comY* by ComX and the effect of the MecA/ClpCP system. Heterogeneity in bacterial protein copy number can be well-described by a physical model of transcription and translation as consecutive stochastic (Poisson) processes, characterized by rates *k*_*r*_ (transcripts per unit time) and *k*_*p*_ (protein copies per transcript per unit time), respectively (Friedman *et al.*, 2006; Taniguchi *et al.*, 2010). In this model the protein copy number *n* in each cell is a random variable drawn from a gamma distribution Γ(*n* | *A, B*). The two parameters *A* and *B* that determine the shape of the distribution are related to *k*_*r*_ and *k*_*p*_, respectively (and to the mRNA and protein lifetimes) (Friedman *et al.*, 2006). Gamma distribution fits to our P*comY* reporter data are shown in Supporting Figure S2.

To model ComX activation of *comY* in the *mecA* deletion mutant (lacking post-translational regulation by MecA/ClpCP), we applied a simple quantitative model in which the P*comX* activity of each cell, as reported by GFP fluorescence, determines the gamma distribution for its P*comY* activity, measured by RFP fluorescence. Specifically, the P*comY-rfp* reporter fluorescence *Y* of a cell is a random number drawn from a gamma distribution Γ(*Y* | *A, B*), for which the parameters are

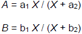

Here, *X* is the P*comX-gfp* reporter fluorescence of that cell. Thus *Y* is directly activated by *X* in a saturating but noisy fashion. We fit this model to a dataset of individual cell RFP and GFP fluorescence values collected on dual reporter (P*comX-gfp*, P*comY-rfp*) Δ*mecA* cells that were supplied with different concentrations of synthetic XIP (defined medium) and then imaged on glass slides. Maximum likelihood analysis gives the four model parameters *a*_1_, *a*_2_, *b*_1_, *b*_2_ for the Δ*mecA* strain as follows: We start with the experimental P*comX* activity measured for each cell, then use the four parameters to define a P*comY* gamma distribution for that cell. We find the probability of that cell’s actual P*comY* activity, given that gamma distribution. The parameter values are then adjusted to maximize the likelihood of the total dataset. (The optimal values are given in Supporting Figure 3.) Given these model parameters we then generate a model simulation for comparison against the data as follows: We use the parameters and the experimental P*comX* activity of each cell to generate its P*comY* gamma distribution, draw a random number from that distribution to obtain a simulated P*comY* activity for the cell, and then plot the resulting simulated P*comY* vs P*comX* values for all cells.

We also tested an alternative model in which environmental XIP concentration, rather than P*comX* activity of a cell, is the determinant of that cell’s P*comY* activity. In this model *X* in the above equations refers to the XIP concentration supplied to a cell. Again, using maximum likelihood, we found the parameters (*a*_1_, *a*_*2*_, *b*_1_, *b*_*2*_) that gave best agreement with the Δ*mecA* data in this alternative model. The scatterplot of Supporting Figure S3, generated by the above simulation procedure, compares the simulated P*comY* to the experimental P*comY* for the Δ*mecA* data.

For the dual-reporter Δ*clpC* mutant, we extended the above model by allowing MecA to sequester, but not degrade ComX. For simplicity we assume that (i) MecA and ComX bind with sufficiently high affinity that a cell can only activate *comY* to the extent that its ComX copy number exceeds its number of MecA copies, leaving some available ComX; ii) The activation of *comY* by the available ComX is as described in the Δ*mecA* model above (and with the same parameters); (iii) the MecA copy number *M* in a cell is a stochastic variable drawn from a gamma distribution Γ(*M* | *A*,*B*) whose *A* and *B* parameters are fixed, independent of XIP concentration. If *X* is the P*comX-gfp* activity of a given cell, then *X’* = *X-M* is the amount of ComX available after sequestration by MecA. Given a GFP measurement of *X* for a cell, the MecA gamma distribution Γ (*M*=*X*-*X*’ | *A*,*B*) determines the probability that *X’* copies of ComX are available to activate *comY*. This *X’* determines the probability distribution for *Y* (the P*comY-rfp* response) by the above model. Averaging over the MecA distribution then gives a prediction for both the average behavior and cell-to-cell variability in the dependence of P*comY-rfp* on P*comX-gfp*, in the presence of MecA.

Taking the P*comY-rfp* activation parameters obtained in the Δ*mecA* fit, we therefore analyzed individual cell P*comX/*P*comY* data that was collected on Δ*clpC* cells that were supplied with different XIP concentrations and imaged on glass slides. We then found the *A* and *B* values for the MecA distribution that maximize the likelihood of the P*comX*/P*comY* dataset, given the sequestration model. Using those parameters, we then generated a simulation of the P*comY* versus P*comX* activity. We compared these results to the experimental data for the Δ*clpC* strain. In plotting the simulation, we modelled the weak red auto-fluorescence background in the data by adding baseline Gaussian noise of 3 ± 0.8 red fluorescence units; this baseline is small compared to the typical red fluorescence (~10^2^-10^4^ units) of *comY* activated cells.

## Acknowledgments

This work was supported by 1R01 DE023339 and T90 DE021990 from the National Institute of Dental and Craniofacial Research.

